# A highly multiplexed assay to monitor pathogenicity, fungicide resistance and gene flow in the fungal wheat pathogen *Zymoseptoria tritici*

**DOI:** 10.1101/2022.07.18.500446

**Authors:** Hadjer Bellah, Gwilherm Gazeau, Sandrine Gélisse, Reda Amezrou, Thierry C. Marcel, Daniel Croll

**Author notes:** Corresponding authors (TM), (DC).

## Abstract

Crop pathogens pose severe risks to global food production due to the rapid rise of resistance to pesticides and host resistance breakdowns. Predicting future risks requires monitoring tools to identify changes in the genetic composition of pathogen populations. Here we report the design of a microfluidics-based amplicon sequencing assay to multiplex 798 loci targeting virulence and fungicide resistance genes, and randomly selected genome-wide markers for the fungal pathogen *Zymoseptoria tritici*. The fungus causes one of the most devastating diseases on wheat showing rapid adaptation to fungicides and host resistance. We optimized the primer design by integrating polymorphism data from 632 genomes of the same species. To test the performance of the assay, we genotyped 192 samples in two replicates. Analysis of the short-read sequence data generated by the assay showed a fairly stable success rate across samples to amplify a large number of loci. The performance was consistent between samples originating from pure genomic DNA as well as material extracted directly from infected wheat leaves. In samples with mixed genotypes, we found that the assay recovers variations in allele frequencies. We also explored the potential of the amplicon assay to recover transposable element insertion polymorphism relevant for fungicide resistance. As a proof-of-concept, we show that the assay recovers the pathogen population structure across French wheat fields. Genomic monitoring of crop pathogens contributes to more sustainable crop protection and yields.

## Introduction

Approximately 30 percent of all crop diseases are caused by fungi [1]. Plant pathogenic fungi affect crops at various life cycle stages and plant tissues, including seeds, root and leaf development, and inflorescence [2–5]. Yield reductions by pathogenic fungi cause food insecurity and economic losses [6,7]. Crop protection is primarily achieved through the application of a variety of fungicides and resistance breeding [8,9]. However, fungal pathogens have evolved resistance to all major fungicides currently in use [10]. In addition, efforts to breed resistant crop varieties have repeatedly been defeated by rapid evolutionary change in pathogen populations allowing them to circumvent resistance mechanisms [9]. Predicting future breakdowns in fungicide efficacy and crop resistance remains challenging. Fungicide resistance is monitored across the European continent by analyzing mutations in known target genes related to the fungicide mode of action [11,12]. However, the rise of pathogen strains defeating crop resistance is not comprehensively monitored. Notable exceptions include the screening of rust fungi [13–16]. Notably, MARPLE (mobile and real-time plant disease) is a genomics-informed monitoring tool developed to quickly detect wheat rust fungal pathogens *in situ* using using Nanopore sequencing [17]. To reduce damage caused by plant pathogens, a timely and accurate detection of both fungicide resistance mutations and mutations associated with the defeat of crop resistance is essential.

Fungal plant pathogen populations that evolved resistance to specific fungicides harbor numerous mutations in or nearby the genes encoding the targets of the chemical compounds [10,18–20]. Similarly, pathogen populations virulent on previously resistant crop varieties have often mutated or deleted a specific set of genes that encode proteins recognized by the plant immune system [21–24]. Fungicide resistance has traditionally been detected using *in vitro* fungicide sensitivity assays [25,26]. Such analyses require the isolation and culturing of individual fungal strains that can then be tested for growth on media containing different fungicide concentrations. The fungicide dose that effectively inhibits growth by 50% is determined for comparison among samples (*i*.*e*., EC50) [25,27]. The method is laborious and limited to fungal species that can be cultured in absence of the host. With advances in molecular techniques, a number of genetic screening methods have been developed including Sanger sequencing, TaqMan assays based on fluorescently-tagged, allele-specific probes [28]. In general, such screening approaches are labor-intensive and have low potential for multiplexing large numbers of individual loci. Virulence surveillance of fungal plant pathogens has been implemented using simple sequence repeat (SSR) markers [29,30] to distinguish the virulent Ug99 race from other *P. graminis f. sp. tritici* lineages [30,31]. However, these SSR makers have been less useful in distinguishing different Ug99 race group members [32]. Besides, virulence monitoring was also performed using loop-mediated isothermal amplification (LAMP), see *e*.*g*. for the wilt *Fusarium oxysporum f. sp. lycopersici* (*Fol*) [33]. However, LAMP assays can be expensive given costs of individual probes.

The advent of next generation sequencing (NGS) approaches has removed a series of limitations in pathogen monitoring. The most general application of NGS techniques is whole genome sequencing (WGS) that can be used to detect single nucleotide polymorphisms (SNPs) and structural variation [34]. Applications of WGS have contributed to the mapping and characterization of virulence and resistance factors primarily through genome-wide association mapping [10,23,35,36]. Low-cost, high-throughput methods based on NGS include reduced representation sequencing genotyping methods such as restriction-site-associated DNA sequencing (RAD-seq) and Genotyping-by-Sequencing (GBS), both methods rely on restriction enzymes to reduce genome size and complexity and exploring SNPs adjacent to restriction enzyme sites [37,38]. However, such genotyping approaches assess only mutations near restriction enzyme cut sites. Applications in fungal pathogens include fine-grained population structure analyses, assessments of recombination rates, mapping of quantitative traits as well as the ability to establish virulence profiles for clonal pathogens [39–44]. The analysis of individual regions involved in fungicide resistance has been improved by the recent development of a PacBio long-read sequencing assay based on the multiplex amplification of target genes in fungal wheat pathogen *Zymoseptoria tritici*. The main advantage is the ability to generate long-reads capturing significant haplotype information of individual strains revealing a series of alterations conferring increased resistance in response to different commercial fungicides. However, due to the varying amplicon sizes generated by this assay two separate multiplex PCRs were required to separate shorter and longer amplicons [26]. High degrees of multiplexing for amplicons and samples were recently achieved using two parallel approaches for animal and plant species. Genotyping-in-thousands by sequencing (GT-seq) is based on multiplex PCR targeted amplicon sequencing to simultaneously genotype thousands of loci and hundreds of samples in a single Illumina sequencing run [45]. A limitation of this approach is the extended time required for its development (about ~4 months according to [46]. One challenge to overcome is imbalanced amplification of individual loci and samples. Such bias can be reduced by the use of Fluidigm microfluidics assays, which physically separate sets of amplicons and samples [47]. The fungal pathogen *Z. tritici* causes one of the economically most important wheat diseases called Septoria tritici blotch (STB) [48]. The pathogen has emerged at the onset of wheat domestication in the Middle East [49] and has since spread to all wheat-producing areas of the world [50]. Populations have evolved resistance to all commercially used fungicides and repeatedly across continents [23]. Major routes to resistances included the rise of mutations in genes encoding the targets of the fungicide, in particular in *CYP51* encoding the target of azoles [8,10]. Furthermore, upregulation of the transporter gene *MFS1* due to the insertion of transposable elements in the promoter region contributed to azole resistance [20]. The rise of succinate dehydrogenase inhibitor (SDHI) resistance mutations are the most recent of the observed gains in resistance (Fungicide Resistance Action Committee, FRAC, 2021). In parallel to the rapid evolution to resist fungicides, *Z. tritici* has also surmounted most known resistance factors segregating among wheat cultivars [51]. Association mapping in *Z. tritici* has recently revealed specific mutations underlying the gain of virulence on previously resistant wheat cultivars including cultivars carrying the resistance gene *Stb6* and others [35,52,53]. Recently, Amezrou et al. (unpublished) identified an additional 58 candidate pathogenicity related genes based on association mapping on 12 wheat differential cultivars. The genes linked to gains of virulence are typically referred to as effector genes and show rapid evolutionary change in populations of *Z. tritici* [22,52,53]. Gene flow among *Z. tritici* populations is leading to significant weak differentiation at the continental scale and high local diversity [50,53,54]. Monitoring of fungicide resistance mutations is mainly achieved through the sequencing of target genes including the recent development of long-read sequencing assays [26]. A joint monitoring of pathogenicity related mutations and genetic diversity is lacking though.

Here, we report the design and validation of a microfluidics based multiplex targeted amplicon sequencing assay that allows the simultaneous monitoring of mutations in fungicide resistance genes and effector genes associated with a wide range of host resistance factors. In addition, we enable the monitoring of hundreds of equally spaced polymorphisms along chromosomes to identify recent changes in the genetic composition of pathogen populations. We validate the performance of the assay using replication, sensitivity analyses to low input DNA, mixed samples as well as the performance on DNA directly obtained from infected wheat leaves.

## Results

### Marker design based on whole-genome sequenced individuals across species

We used whole-genome sequencing datasets of 632 *Z. tritici* isolates collected in Oceania (Australia, New Zealand), the United States, Switzerland, France, and Israel to identify segregating SNPs and improve the design of a total of _79_8 amplicons of ~200 bp of length (except for the MFS1 and *ZtSDHC3* loci). The short and largely identical amplicon lengths improve PCR efficiency and balance among loci. Known polymorphism within the species was used to mask sites to avoid primer mismatches and amplification drop-outs (Fig. 1A). We designed 25 amplicons across genes associated with fungicide resistance including *CYP51*, alternative oxidase (*AOX*), beta-tubulin (*TUB1*), *SDH1-4* genes including *ZtSDHC3*, as well as *cytochrome b (CYTB)* (Table B in the File S1). For each gene, we prioritized amplicons covering non-synonymous substitution if available. Due to the complexity of the transposable element insertion polymorphism in the promoter region of the transporter gene *MFS1*, we designed a total of 16 primer pairs for amplicons matching known sequence variants near three insertion sites [20] (Table B in File S1). For loci associated with pathogenicity on diverse cultivars, we retained a set of 67 amplicons successfully passing primer design (Table B in File S1). We also randomly selected SNPs at ~50 kb distances to monitor the genetic make-up of populations for a total of 691 designed amplicons across all chromosomes (Table B in File S1). The random SNP set also included by chance the previously selected fungicide resistance gene *cytochrome b* (*CYTB*).

**Figure 1:**
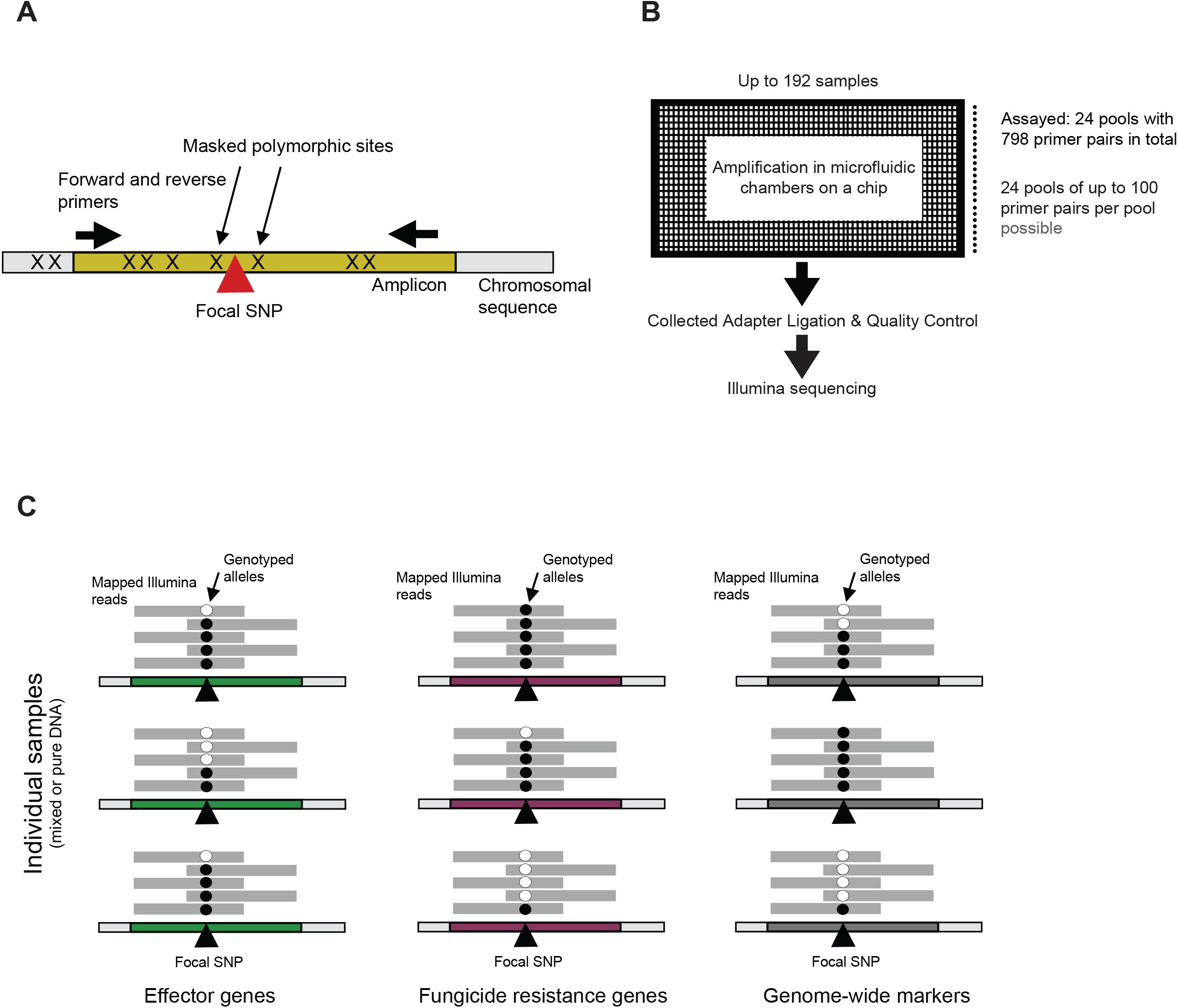
Schematic overview of the targeted amplicon assay design. A) Design of individual amplicons (~200 bp) with primers designed to not overlap known polymorphic sites. B) Schematic overview of the microfluidic chambers of a Fluidigm Juno chip accommodating up to 192 samples and 24 pools of primers (each up to 100 primer pairs). Following amplification in microfluidic wells, barcoded products are pooled and finalized for Illumina sequencing. C) Genotypes of individual samples (pure or mixed individuals) are assessed by analyzing mapped reads at each locus in the genome. Markers were designed for three different categories including effector genes, genes encoding targets of fungicides and genome-wide evenly spaced markers.

### Assessment of loci quality across the targeted sequencing assay

We performed targeted sequencing of all _79_8 loci based on the Fluidigm Juno system in a single run using microfluidics (Fig. 1B). The 192 samples included four sets of pure DNA from different isolates mixed in equal proportions, ten samples including each DNA of the same three isolates in different proportions, and 178 samples constituted from extracted leaf material from different wheat fields across France mostly (*i*.*e. n* = 172), Belgium, Ireland and the United Kingdom. The complete set of samples was replicated once for the amplification and Illumina sequencing step. The total sequencing output over both replicates was 2,418,905,407 read pairs and 338.89 Gb. For 31 samples, the amplification and Illumina sequencing procedures failed in either one of the two replicates of each sample, therefore the failed replicates were eliminated. Across a replicate run (*i*.*e*. FC2), samples produced between 5,976-173,049,611 read pairs with numbers broadly consistent between the two replicate runs (Fig. 2A). We found that the mapping rate against the reference genome ranged from 96.93-100% among most sample replicates (Fig. 2B).

**Figure 2:**
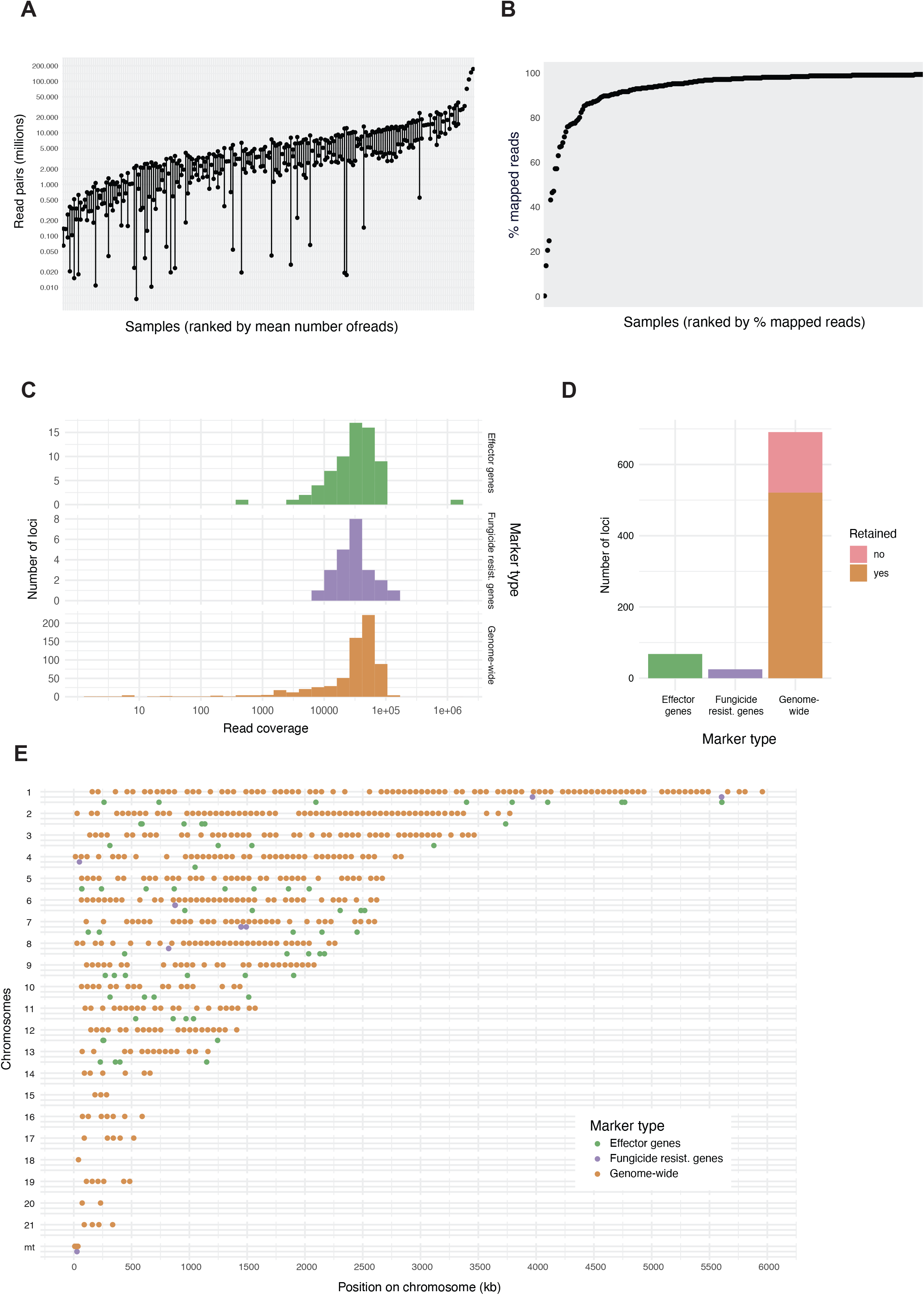
Sequencing data recovered for the amplicon assay and loci assessment. A) Read pairs recovered per sample and replicate. Each sample was amplified and sequenced two times (two different microfluidic flow cells). B) Ranking of percent mapped reads to the reference genome per sample (including both replicates if available). C) Number of reads mapped per locus for the three different categories of markers. The read numbers correspond to the total obtained from four pooled samples performed in replicates. D) Summary of loci retained after read number filtering. Only genome-wide markers were removed if they failed filtering criteria. E) Overview of retained markers per category across the 21 chromosomes and mitochondrion.

To assess the faithful amplification of individual loci, we first focused on the four samples with mixtures of pure fungal DNA of 26 to 30 isolates. Combining the two replicates, we used eight samples to evaluate sequencing read coverage across the 782 amplicons designed outside of the *MFS1* region. We found 17 loci with a read depth of 0. The highest read depth was 1,779,927 for an effector locus on chromosome 1. For the set of genome-wide, equally spaced amplicons on core chromosomes, we accepted the locus if the read counts were between 20,000 and 100,000 in the retained samples (Fig. 2C). We considered this read count range to reflect the loci consistently amplifying across samples and not showing evidence for duplications. With this filter, we discarded 149 loci falling outside of the read count range (Fig. 2D). For randomly selected markers on accessory chromosomes, we expected lower amplification success because not all isolates of the species carry the locus. We retained loci with a read count between 10,000 and 100,000 in the set of reference samples leading to the rejection of 21 loci (Fig. 2C-D). For randomly selected mitochondrial markers, we found read counts ranging from 203,502 to 1,372,965 in the set of reference samples reflecting the high copy number of mitochondria compared to the nuclear genome. All 12 randomly selected mitochondrial loci were kept. For effector loci, the number of mapped reads ranged from 502 to 1,779,927 reads indicating significant variation in the amplification success and possibly copy number (Fig. 2C). We retained all 67 designed amplicons due to the general interest in polymorphism at such loci (Fig. 2D). For resistance gene loci, the number of mapped reads ranged from 2,986-1,372,965 reads (Fig. 2C). As for effector gene loci, all 24 designed amplicons were retained (Fig. 2D). In addition, we retained the amplicon for the mitochondrial resistance locus of *CYTB* with a read count of 1,372,965. In summary, we retained 521 high-quality loci representing 75% of the randomly selected markers designed for genetic structure analyses, as well as all 67 effector and 24 fungicide resistance loci (Fig. 2D).

### Reproducibility among replicate assays and recovery of allele frequencies

To assess the reproducibility of the sequencing assay, we repeated the amplification and sequencing procedure two times. We found that the number of read pairs recovered for each sample were positively correlated between replicates (*r* = 0.78, p-value < 0.0001; Fig. 3A). We also found a positive correlation in the mapping rate of reads recovered from the same samples (*r* = 0.85, p-value < 0.0001; Fig. 3B). To investigate effects on allele frequencies assessed for mixed samples, we compared the pooled DNA of population 41 sample. We used allele frequencies estimated from read depth for the reference and alternative allele at SNP loci. Reference allele frequencies at 201 SNP loci calculated in both replicates of each sample were highly correlated (*r* = 0.89, p-value < 0.0001) with outliers corresponding to poorly covered loci in either one of the two replicates of the same sample (Fig. 3C).

**Figure 3:**
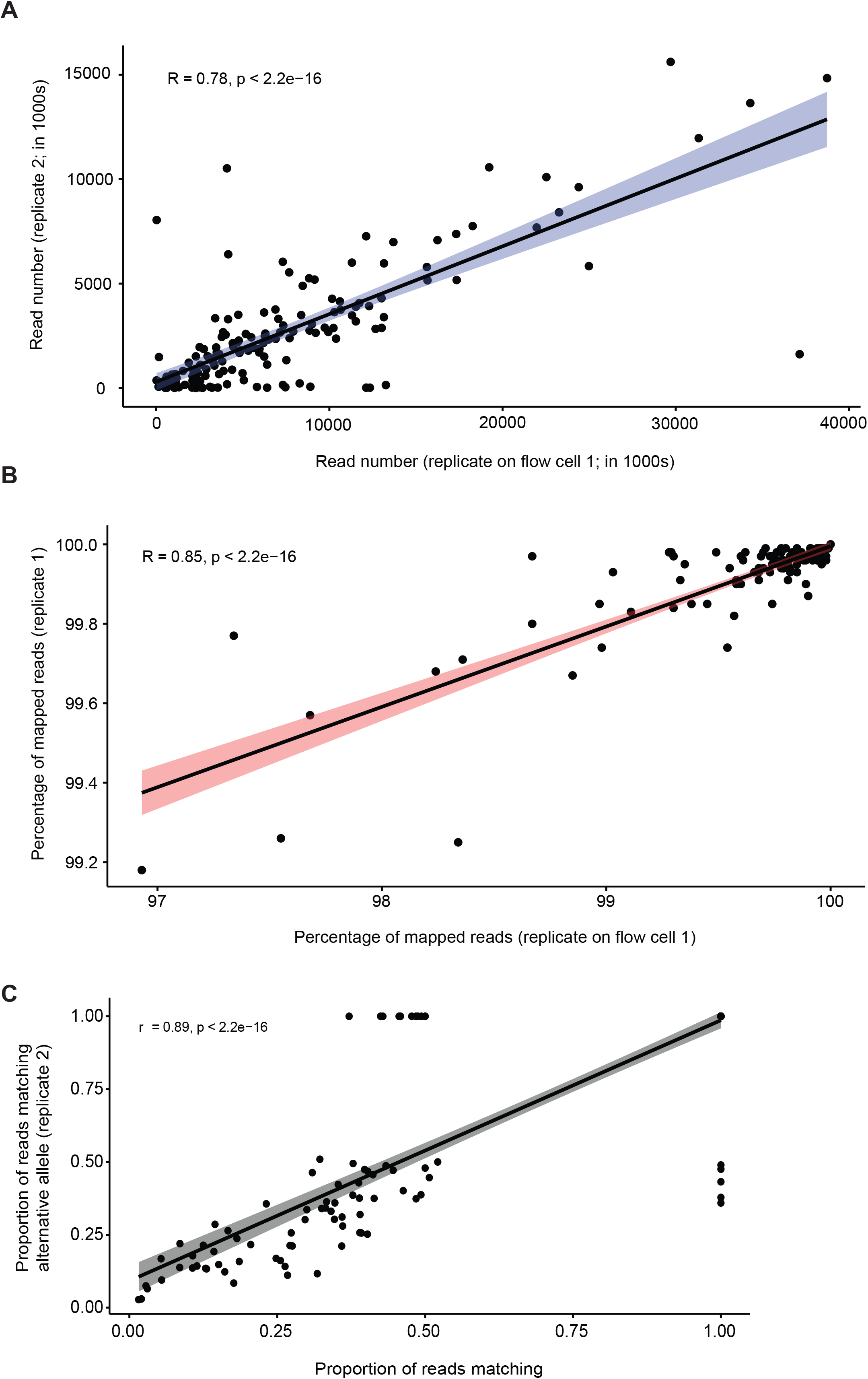
Consistency between replicate runs of the amplicon assay. A) Read numbers per sample and B) percentage of reads mapped to the reference genome. C) Comparison of alternative allele frequencies within samples between the two replicate runs for each sample.

Furthermore, we analyzed allele frequencies in ten samples (*i*.*e*. G1-G10) constituted from a mix of pure DNA from the same three isolates in different proportions (INRA10-FS1006, INRA10-FS1022, IPO-09455; Fig. 4; Table A in Supplementary File S1). We used existing whole genome sequencing and SNP calling data for the three isolates to assess polymorphism across the genome [55]. Using the known dilutions of pure DNA, we established the expected frequencies of reference alleles (*i*.*e*. matching the allele present in the reference genome IPO323) or alternative alleles across loci. Then, we analyzed mapped reads from the targeted sequencing assay from the mixed samples G1-G10 across all amplicons to identify the proportion of reads matching the reference allele (Fig. 4). If the targeted sequencing assay faithfully amplified DNA in mixed samples, the expected reference allele frequency in the mixed samples should match the recovered proportion of reads matching the reference allele. Across the ten different mixed samples, the match in reference allele frequencies was high in most samples (linear regression with *R*^*2*^ > 0.55 in 7 out of 10 mixtures). The mixed sample G1 showed no association between DNA dilutions and recovered allele frequencies and two additional samples (G5 and G6) showed weak associations (*R*^*2*^ = 0.27-0.45).

**Figure 4:**
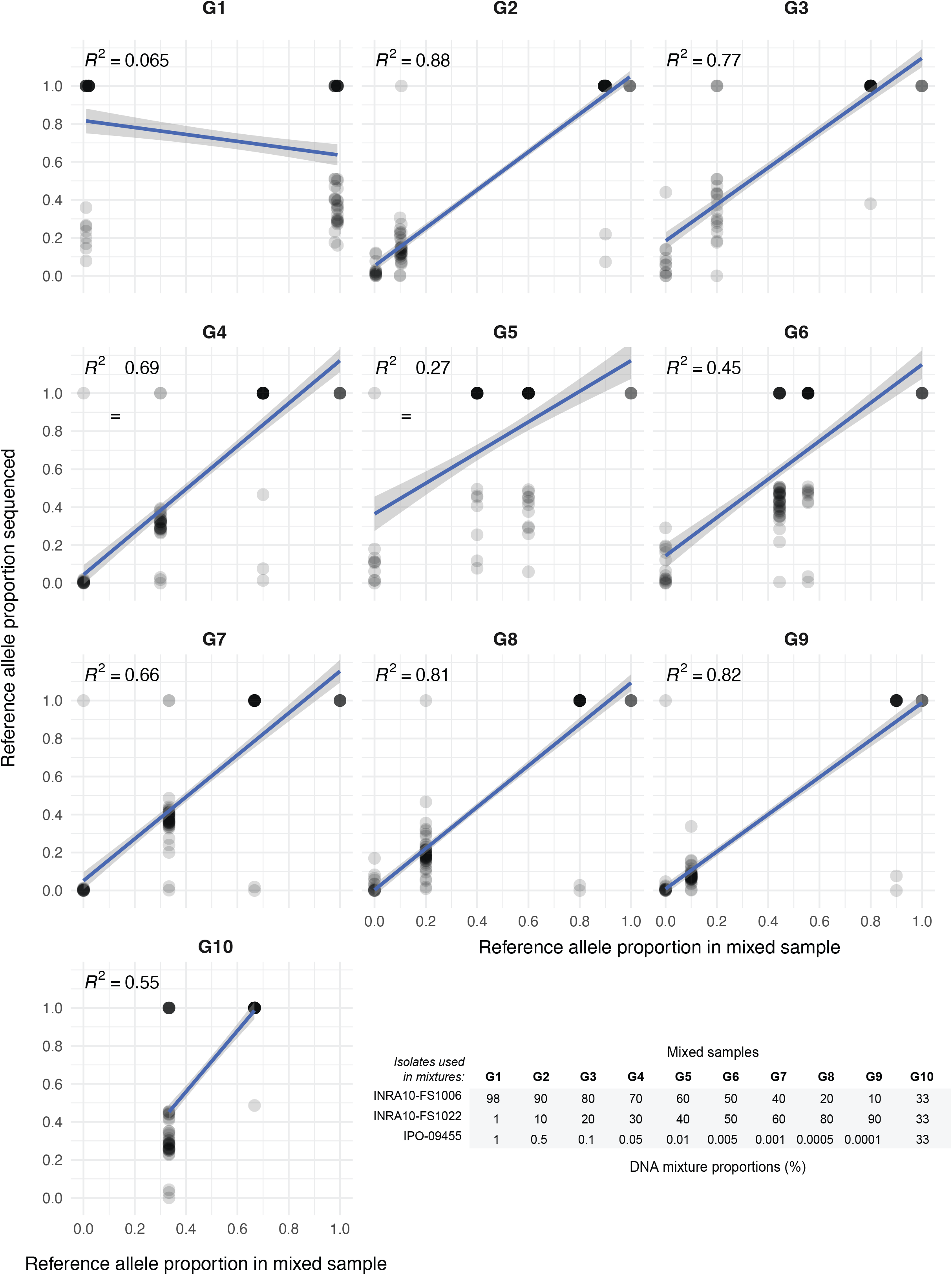
Evaluation of mixed sample analyses. Ten samples (G1-G10) contained mixed DNA of three different isolates (INRA10-FS1006, INRA10-FS1022, IPO-09455) varying in proportions. Genotypes of each of the isolates were retrieved from whole-genome sequencing of pure isolates and assigned as reference alleles (*i*.*e*. matching the allele present in the reference genome IPO323) or alternative alleles. Using known genotypes of the three isolates, reference allele proportions were defined according to the dilutions in mixed samples G1-G10. Amplicon sequencing data of mixed samples was screened for all genotyped SNPs to assess the proportion of the reference allele among all mapped Illumina reads. Only SNPs with a minimum read coverage of 50 were used. Regression *R*^*2*^ were calculated based on a linear model.

### SNP monitoring in fungicide resistance genes

We investigated the amplification success for the amplicons covering the *CYP51* locus using eight sets of mixed pure fungal DNA samples with read counts ranging from 10,115 to 31,323 reads. The genotyping of infected wheat leaf samples from the field revealed that the target SNPs were indeed polymorphic. For *CYP51* and the other fungicide resistance associated genes such as *TUB1, AOX, SDH2* and *SDH3* the dominant genotype per wheat leaf varied among samples (see File S1, Tables D and E). The reference genome isolate IPO323 is generally susceptible to different fungicide classes. Hence, the allele carried by the reference genome is likely associated with higher susceptibility. Consistent with recent gains in fungicide resistance, mutations in the beta-tubulin and *CYP51* locus tended to be different from the reference genome (*i*.*e*. the alternative allele, Table E in File S1). Loci without recent strong recent gains more likely retained the IPO323 genotype (*i*.*e*. reference allele, Table E in File S1).

### Amplicons for the promoter region of MFS1

The amplicons designed for the promoter region of *MFS1* are matching known haplotypes differing in their insertion of transposable element sequences. Due to the sequence complexity, we chose to first cluster sequencing reads into individual amplicons instead of directly mapping reads to a *MFS1* haplotype. Analyzing the 10 samples with different DNA mixtures of three isolates including replicates, we identified 10 sequence clusters with at least 22 reads (lowest number observed in sample G3). We used BLAST to retrieve the subset (*n* =10) of the clustered sequences matching the *MFS1* promoter region. The sequences matched positions from 1-4946 bp (for sample G9) on the consensus *MFS1* sequence with all being upstream of the coding sequence as expected (Fig. 5A). We did not recover any amplicon matching forward and/or reverse primer positions based on the amplicon design (Fig. 5B). However, all amplicons did not match the expected amplicon length most likely due to the complexity of the underlying sequence. Furthermore, the pooled amplification of multiple primer pairs matching the promoter region has likely produced chimeric amplicons in some contexts. We used the retrieved amplicons matching the promoter region to form clusters of near identical BLAST matches based on alignment length and positions. We identified 10 well supported amplicon clusters showing variation in abundance among the analyzed samples (See Table C in File S1).

**Figure 5:**
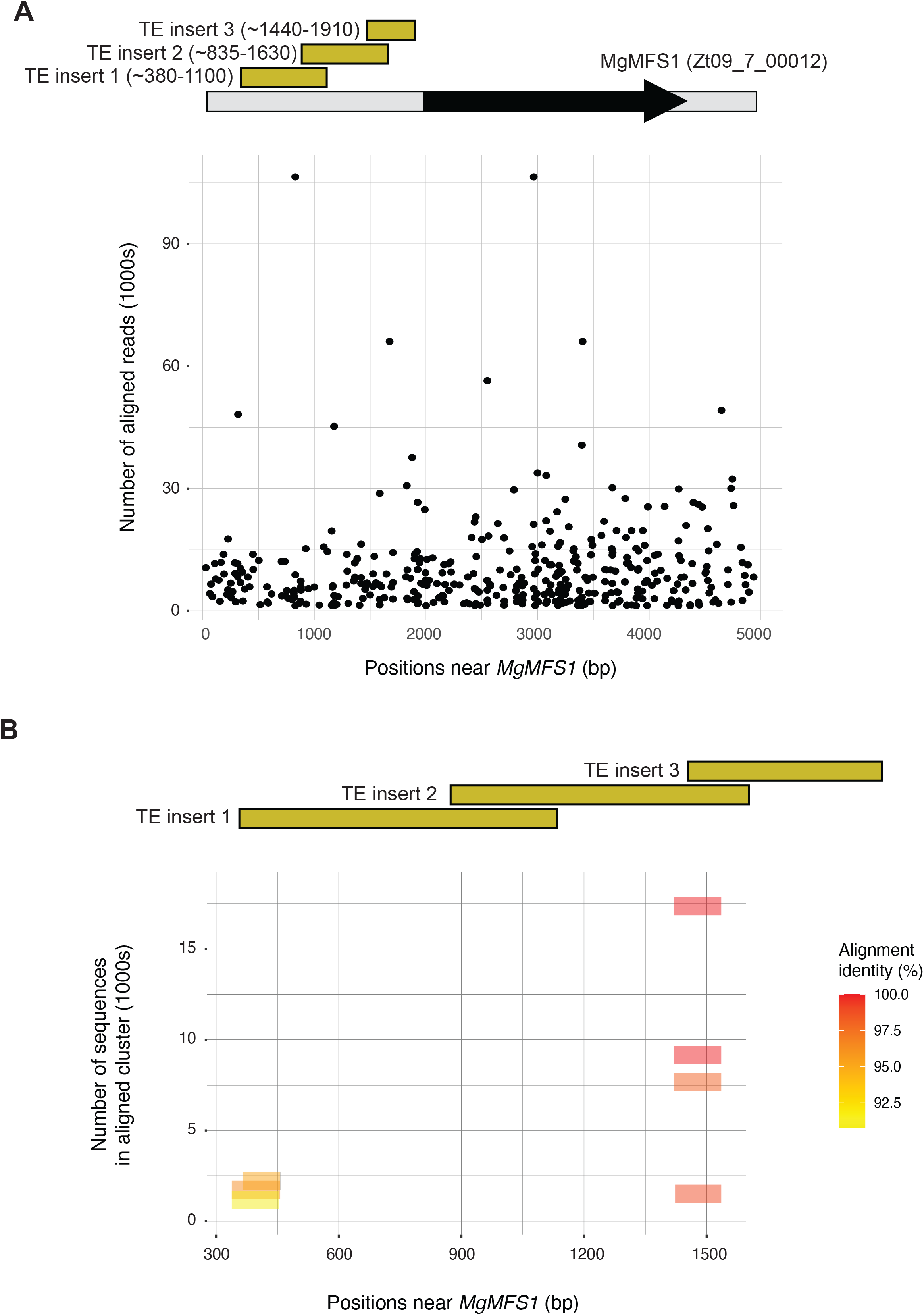
Analyses of amplicons designed on polymorphic transposable element insertions upstream of the multidrug transporter gene *MgMFS1*. A) Overview of the location of amplicons designed for each of three transposable element insertion site (1-3). Multiple amplicons were designed for each insertion site. The aligned reads are shown for positions near the coding sequence of *MgMFS1* for sample G9 (only positions with >10 reads mapping are shown). B) After read clustering for sample G9, consensus sequences were blasted against positions near the coding sequence of *MgMFS1*. The horizontal bars indicate the extent of a BLASTn alignment with colors indicating the percent identity of the alignment. The vertical position indicates the number of sequences that were clustered for the aligned consensus sequence.

### Genetic differentiation in French and European wheat field populations

We used the 158 wheat leaf samples infected by *Z. tritici* collected from fields across France with at least five genotyped samples per location and additional samples from Belgium, Ireland and the United Kingdom to assess the genetic structure using the genome-wide marker set (Supplementary Figure 1 in File S2). Based on a principal component analysis of 85 genome-wide SNPs, we found no clear differentiation among samples originating from different countries (Fig. 6A). Focusing on the genetic differentiation among French regions (*n* = 82 genome-wide SNPs for the French populations only), we found some modest differentiation of genotypes from wheat fields in Midi-Pyrénées and Champagne (Fig. 6B). However, the overall differentiation of the field samples was low with the first and second principal component explaining only ~4%.

**Figure 6:**
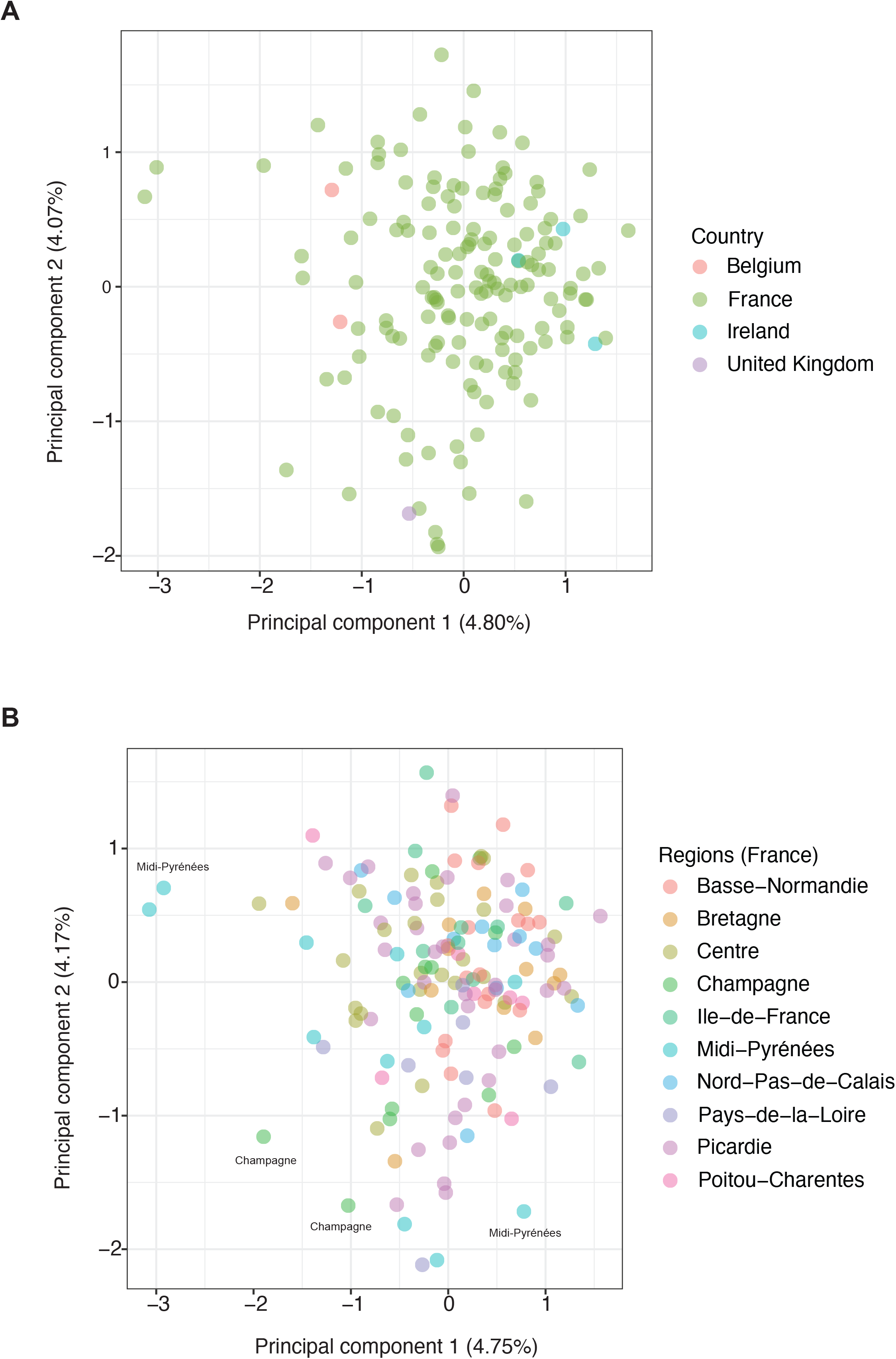
Population structure analyses based on genome-wide markers genotyped on leaf-extracted assemblies of *Zymoseptoria tritici* strains. A) Principal component analysis of wheat leaf samples collected in France, Belgium, Ireland and the United Kingdom and B) the subset of wheat leaf samples collected in France colored by region.

## Discussion

We developed a microfluidics-based amplicon sequencing assay combining the advantages of high-throughput sequencing and multiplex PCR. We assessed the performance of 798 loci to reliably and sensitively genotype randomly selected genome-wide markers, as well as pathogenicity and fungicide resistance-related genes in a diverse set of *Z. tritici* samples. We show that a large portion of the designed markers can be amplified consistently across samples, used to monitor the emergence of relevant mutations and provide an assessment of allele frequencies in mixed samples. The set of genome-wide markers provides means to assess the genetic structure of the pathogen directly from field collected wheat leaves.

Within-species polymorphism can lead to amplification failures due to mismatching primers. We considered this issue particularly relevant for the wheat pathogen *Z. tritici* as the species harbors genetically highly diverse populations within single fields [53]. As expected, we detected a high number of SNPs in regions intended for amplicon design leading to the rejection of amplicon candidates prior to the genotyping stage. Furthermore, we noticed targeted regions with weak amplification success. The poor performance of some primer pairs is most likely explained by a combination of factors. First, we ignored low-frequency SNPs at the masking stage to be able to proceed to the amplicon design for more loci. Second, our species-wide genomic survey of SNPs may have missed polymorphisms present in the assayed samples. The filtering thresholds can be adjusted and more genome sequencing datasets could be included in future amplicon design efforts. Despite some failed attempts at amplifying individual loci, we obtained high degrees of sequencing read coverage for most loci. Most samples yielded hundreds to thousands of reads for each locus. Such deep coverage across the amplicon assay provides a detailed picture of genotypic diversity particularly for mixed samples directly obtained from infected leaves. A major limitation with the multiplexed amplicon sequencing assay is the shortness of the amplified sequence (~200 bp). The short amplicon length ensures a high degree of multiplexing by providing stable amplification across the entire assay. However, longer amplicons would be needed to recover entire sequences (*i*.*e*. haplotypes) of the azole resistance locus *CYP51* or several effector genes of interest. A potential solution would be to design overlapping amplicons to cover an entire locus. However, this approach was unsuccessful *e*.*g*. for the effector gene *AvrStb6* providing no sufficiently conserved sections inside or adjacent to the coding sequence for an overlapping amplicon design. Limitations in amplicon length and haplotype resolution can be overcome using long-read sequencing as developed to monitor fungicide resistance loci in *Z. tritici* [26]. Long-read sequencing may also help to overcome issues with amplifying the highly polymorphic promoter region of *MFS1*. Long amplicons could capture the entire promoter region instead of focusing on individual insertion points. However, long-read approaches are not well-suited to amplify hundreds of loci consistently across many samples. Ultimately, a combination of different approaches performing highly multiplexed short reads sequencing and separate long-read sequencing for the most complex loci will be required.

A versatile pathogen genotyping assay should perform well with low-input pure fungal DNA as well as mixed samples containing DNA both from multiple strain genotypes and the host (*i*.*e*. wheat plants). We find that the assay replicated well across most of the tested sample types both in terms of the number of recovered reads per sample as well as the proportion of reads that could be mapped to the *Z. tritici* reference genome. Besides, we found that in mixed samples (*i*.*e*. containing more than one genotype), the assay reproduces well the allele frequencies across the two independent genotyping runs. We also assessed the ability of the assay to recover allele frequencies of mixtures of known isolates. Using known genotypes of three isolates as a control, the amplicon assay recovered well the allele frequencies in most tested mixtures. The weak performance of some individual mixtures is likely due to errors during handling rather than a general issue of reproducing allele frequencies. The accurate recovery of allele frequencies in a mixed sample is clearly contingent on sufficient sequencing depth though and we have evaluated the performance only at loci with ≥50 mapped reads. The genotyping of transposable element insertions in the *MFS1* promoter region was not conclusive. The overlapping amplicons and very high levels of sequence polymorphism prevented a clear assignment of amplicons to transposable element insertion genotypes, but our data opens up a path for a more comprehensive design strategy to capture inserted sequences.

The microfluidics-based multiplex PCR targeted amplicon sequencing requires bioinformatics analyses both for the design of the amplicons and for the genotyping after a successful run. Nearly all designed amplicons and associated primer sequences could be used also outside of a microfluidics protocol. A technically less demanding version of our approach is typically referred to as GT-seq, which consists in amplifying loci in large pools of primer pairs and indices to distinguish samples [45]. Given the short amplicons, using individual primer pairs for targeted qPCR assays would also be possible. The number of recovered loci for targeted amplicon sequencing remains below untargeted approaches such as RAD-seq and GBS. Untargeted reduced-representation approaches provide however only genome-wide information on genetic differentiation. This may be informative *e*.*g*. for virulence profiles in clonal pathogens [43], however this approach is unsuitable to recover genotypes at specific loci. Targeted amplification such as the microfluidics based multiplex PCR performs also well in mixed samples. RAD-seq and GBS are unlikely to perform well if substantial proportions of plant DNA are present, because large plant genomes will typically contain many more restriction sites compared to fungal genomes.

The developed microfluidics-based targeted amplicon assay allows a cost-effective and reproducible monitoring of hundreds of loci to track mutations at pathogenicity loci and fungicide resistance evolution in field populations. The integration of genome-wide markers greatly enhances the quality of pathogen monitoring by providing information about patterns of gene flow. Our study revealed only weak differentiation across Western European countries and among French regions consistent with high levels of gene flow and genetic diversity [55]. Knowledge of genetic structure can help identify recent movements of the pathogen due to natural or human-mediated dispersal. The rapid rise in resistance of *Z. tritici* populations after the application of fungicides can more effectively be monitored due to the large number of loci that can be assayed simultaneously. Furthermore, tracking mutations at effector loci opens new opportunities to track adaptation to different wheat cultivars across regions. With the availability of whole genome sequencing data for an increasing number of crop pathogens, the targeted amplicons could be expanded to simultaneously or separately genotype other major pathogens including rusts to improve the surveillance and management of crop diseases globally.

## Methods

### Genome sequences used for the design of the assay

The amplicons were designed based on known polymorphisms within the species. Polymorphic sites were used both to select SNPs to amplify but also to mask polymorphisms to avoid primer binding mismatches. For this, we used whole genome sequencing information from 632 *Z. tritici* isolates collected across the global distribution range of wheat. Isolates included six different populations with a sample size of 29-178. A total of 88 isolates were collected in Australia including Tasmania in 2001 and 2015 [22]. Additional isolates from Oceania included 75 isolates collected in New Zealand in 2013 and 2015 [55]. A total of 154 isolates were collected in Oregon, USA, in 1990 and 2015 [22]. 178 isolates were in wheat fields near Zurich in Switzerland in 1999 and 2016 [22,53] and 29 isolates were isolated in the Nahal Oz region in Israel in 1992 [22]. Finally, 108 isolates were retrieved from a panel of French isolates [35].

### SNP calling and identification of polymorphisms for the amplicon design

We performed read alignment and SNP discovery for the generated genomic datasets, as previously described [22,35]. In summary, we trimmed raw Illumina reads using Trimmomatic v. 0.38 [56] and mapped retained reads to the reference genome IPO323 [57] using bowtie v2.3.5 [58]. We used the Genome Analysis Toolkit (GATK) v4.0.1 [59] including the HaplotypeCaller tool to identify candidate SNPs. We filtered for a set of high-quality polymorphisms using the GATK VariantFiltration tool and vcftools v.0.1.15 [60]. A more extensive description of the filtering procedures and validations are available [61].

### Polymorphism selection for neutral markers, pathogenicity and fungicide resistance genes

Effector candidate genes were retrieved from GWAS focused to identify candidate effectors interacting with major wheat resistance genes (Amezrou et al., unpublished)[52,61]. We included 65 candidate effector genes showing a significant association for symptom development on at least one wheat cultivar. We designed at least one amplicon overlapping the most significantly associated SNP in each of the effector genes. If a significantly associated SNP could not be reproduced in the worldwide isolate collection, a random nearby SNP (within ~200 bp) was selected as the target for the amplicon design. If a different SNP was selected, we filtered for SNPs with a minor allele count of 5 and a minimal genotyping rate of 80%. For the effector gene *AvrStb6*, we designed two additional amplicons to cover polymorphism in the coding sequence. To monitor fungicide resistance gene mutations, we covered 25 genes related to fungicide resistance in *Z. tritici* populations including the mitochondrial genes *CYTB* and *AOX*, the nuclear genes beta tubulin 1 (carbendazim resistance), *CYP51* (azole resistance), as well as *SDH1, SDH2, SDH3* and *SDH4* (SDHI resistance). The amplicons covered resistance mutations if known for the species. If no mutation was previously documented in *Z. tritici*, the amplicon covered randomly selected SNPs in the coding sequence. Similar to the procedure for effector loci, if a known SNP associated with fungicide resistance could not be recovered, we selected a SNP within ~100 bp (minor allele count of 3, minimum genotyping rate 80%). The broader inclusion of polymorphisms for filtering was possible due to the generally lower degree of detected variants in resistance genes. We defined an additional amplicon to target the paralog of *SDH3* (*ZtSDHC3*) [62]. For this, we analyzed the paralog sequence discovered in the pangenome of *Z. tritici* [63].

Multidrug fungicide resistance in *Z. tritici* is mediated by transposable element insertions in the promotor region of the transporter *MFS1*. We designed 16 amplicons covering three previously reported transposable element insertions and haplotypes [20]. The amplicons were designed to either amplify if an insertion was present or not. Amplicons were designed on a consensus sequence of previously described haplotypes [20]. In addition to polymorphisms related to pathogenicity and fungicide resistance, we randomly selected equally spaced polymorphisms along all 21 chromosomes to capture neutral population structure. For this, we selected 691 SNPs with a minor allele frequency of 5% and a minimal genotyping rate of 80%. SNPs were selected at a distance of 50 kb (if available) using the --*thin* option in *vcftools*. In summary, a total of _79_8 amplicons were designed for pathogenicity, fungicide resistance as well as gene flow tracking across *Z*.*tritici* populations. See Table B in File S1 for details on all selected effector and fungicide resistance genes as well as whole genome neutral markers.

### Amplicon design

For genome-wide markers and markers in effector and fungicide resistance genes (except *ZtSDHC3* and MFS1), we extracted a 401 bp sequence from the reference genome centered on the SNP to target. The extracted sequence was centered around the target SNP, which was marked by IUPAC code and parentheses according to company instructions. The sequence was then used to define primers amplifying a ~200 bp stretch of DNA including the target SNP. The amplicon length was limited to ~200 bp to ensure efficient and balanced amplification across loci. To improve amplification success across a broad range of *Z. tritici* genotypes, we masked known polymorphic sites on the sequence containing the targeted SNP to prevent accidental primer design in known polymorphic regions. We used bcftools v1.9 [64] to mask non-target sites showing evidence for polymorphism in the panel of 632 analyzed isolates using the -I option of the consensus command and re-wrote sequences with samtools v1.9 [65]. For resistance and pathogenicity loci, we used a minor allele count of 3 and 5, respectively to consider the polymorphism for masking. For genome-wide markers, we used a minor allele frequency cut-off of 5%. If the resulting sequence contained more than 10% masked sites, the amplicon was not considered further. Additional sequences were excluded by Fluidigm Inc. if the sequences failed to yield adequate primer candidates for the desired ~200 bp amplicons. If the initial amplicon design had failed, we repeated the procedure for effector loci but relaxed the filter to consider only SNPs with a minor allele count of ≥25.

### Samples included for the validation of the amplicon sequencing assay

We assessed the performance of the microfluidics assay using different sets of samples collected from wheat fields in Europe. Four samples included equimolar DNA mixtures of 26 to 30 isolates obtained by culturing single spore isolates from field-collected wheat leaves. Three single spore isolates identified as INRA10-FS1006, INRA10-FS1022 and IPO-09455 were collected in 2009 and 2010 in the Ile-de-France region and were used to create DNA mixtures in ten different proportions (samples G1-G10). Finally, 178 samples were obtained by extracting DNA directly from infected wheat leaves collected in different regions of France, Belgium, Ireland and United Kingdom (Table A in File S1). No permit is required to collect naturally infected wheat leaves.

### DNA extractions and microfluidics assay

DNA extractions to test the microfluidics assay were performed using the following procedures. For pure cultures and directly from infected wheat leaves using DNeasy® Plant Mini Kit (Qiagen, Hilden, Germany). DNA was quantified using a Qubit 2.0 fluorometer (Thermo Fisher, Waltham, Massachusetts, USA). We followed the Fluidigm Inc. (San Francisco, California, USA) Juno™ targeted amplicon sequencing protocol according to the manufacturer’s protocol. As input DNA, we used the following 1.5-200 ng of total amount (See Table A in Supplementary File S1). We performed the entire microfluidics procedure twice independently on different Juno LP 192.24 integrated fluidic circuits plate (IFC). Libraries were prepared following the manufacturer’s protocol. Target amplicons were generated for each sample and pools of primers using PCR on a specialized thermocycler (Juno system; Fluidigm). Illumina sequencing was performed in paired-end mode to generate 100 bp reads on the NovaSeq™6000 platform at Integragen Inc. (Evry, France) and produced 363.89 Gb of raw sequencing data for both independent chips combined.

### Amplicon sequence data analyses

We used Trimmomatic v0.38 [56] with the following parameters: LEADING:3 TRAILING:3 SLIDINGWINDOW:4:15 MINLEN:36. Due to the short amplicon length compared to the read lengths, we used FLASH v1.2.11 [66] to merge forward and reverse reads per pair into single pseudo-reads. Finally, pseudo-reads were aligned to the IPO323 reference genome using bowtie2 v2.3.5 [57,58]. We assessed individual read counts at each analysis step using MultiQC v.1.7 [67]. After individual genotyping using the GATK HaplotypeCaller tool, we performed multi-sample genotype calling using CombineGVCFs and GenotypeGVCFs [68]. Variant sites were removed if these met the following conditions: QD < 5, MQ < 20, -2 > ReadPosRankSum > 2, -2 > MQRankSum > 2, -2 > BaseQRankSum > 2.

The DNA mixtures (G1-G10) contained three isolates INRA10-FS1006, INRA10-FS1022 and IPO-09455 with existing SNP genotyping information [55]. Isolates in mixed samples were diluted in different proportions to cover a range of isolate mixtures. To assess the reproducibility of allele frequencies of the mixed DNA samples, we analyzed mapped reads at each SNP genotyped using the amplicon sequencing assay. Expected proportions of reference alleles (matching the reference genome IPO323) were inferred in mixed samples using the known genotypes of the isolates. Only amplicon sequencing loci with a minimum read coverage of 50 were considered to reduce noise in allele frequency assessments. For amplicons targeting the promoter region of *MFS1*, we first used seqtk [69] to subsample 10.000.000 reads from large merged paired-end reads FASTQ files and we then performed a clustering analysis of Illumina reads to obtain read sets originating from the same locus. We used CD-HIT-EST [70] with an identity threshold set to 100% to cluster sequencing reads. For each cluster, the representative sequence identified by CD-HIT-EST was aligned to the *MFS1* promoter consensus sequence using BLASTn 2.12.0 [71]. Only BLASTn best hits with a bit score above 100 and identity > 90% were kept. To identify clusters of nearly identical hits based on position and identity, we performed *k*-means clustering with the R packages {factoextra}[72], {clustertend} [73], {cluster} [74], {NbClust} [75]. For each sample, we identified the optimal number of clusters (*K* = 1–10) by performing a silhouette analysis [76].

### Data visualization and population genetic analyses

Data analyses were performed using R 4.0.4 [77]. The R packages included in {tidyverse} [78] were used for summarizing and plotting coverage across loci, visualizing retained SNPs, the outcomes of different filtering stages and genotyping. We used bcftools v1.9 [64] to calculate allele frequencies at SNP loci. The allele frequency correlation between both flow cells chips was analyzed with the R package {report} [79] and visualized using {ggpubr} [80] and {ggplot2} [81]. To analyze genetic diversity and population structure, we performed a principal component analysis (PCA) using the R packages {vcfR} [82], {adegenet}[83], {ade4} [84] and {ggplot2} [81]. For population analyses, we focused only on the second replicate (flow cell) and genome-wide SNPs without effector and resistance gene loci to reflect neutral population structure. Loci were filtered for a minor allele frequency of 0.05 and allowing for 20% missing data (--max-missing 0.8).

## Data availability

Raw sequencing data is available on the NCBI Sequence Read Archive (SRA) under BioProject PRJNA847707 (https://www.ncbi.nlm.nih.gov/bioproject/ PRJNA847707).

## Acknowledgements

We thank Anne-Sophie Walker for providing infected leaf samples that were used to develop the assay. We are grateful for the sequence alignment shared by Sabine Fillinger. The microfluidics assay was conducted on the genotyping platform GENTYANE at INRAE Clermont-Ferrand (https://gentyane.clermont.inrae.fr/), with the help of Rachel Fourdin and Lydia Jaffrelo.

## Author contributions

HB, GG and SG performed analyses, RA and DC provided datasets, TCM provided samples, TCM and DC supervised the work. HB and DC wrote the manuscript with input from co-authors.

## Supporting Information files

**File S1: Supplementary Tables A-E**.

Table A: Samples and sample mixtures included in the microfluidics assay.

Table B: Designed amplicons targeting neutral markers, fungicide resistance and effector genes. Loci check: coverage based assessment of amplification success (see methods). If an originally targeted locus was not recovered in the species-wide SNP call set used for the amplicon design, a nearby SNP was chosen (see last columns for newly selected loci).

Table C: Clustering of reads using CD-HIT-EST followed by mapping to the MFS1 promoter region. Similar blast hits were grouped into K-means based clusters.

Table D: Dominant genotype recovered for wheat leaf samples at fungicide resistance loci. The reference allele refers to the allele known from the reference genome isolate IPO323. Sample genotypes are given as 1 and 0 for reference and alternative allele, respectively.

Table E: Dominant genotype recovered for wheat field samples across fungicide resistance loci. The reference allele refers to the allele known from the reference genome isolate IPO323.

**File S2: Supplementary Figure 1**.

Figure 1: Wheat leaf samples collected in France, Belgium, Ireland and the United Kingdom separated by the cultivar of origin or unknown cultivar (“NA”). See File S1 (Table A) for details on the sample origins.

