## Supplementary Figure for "A highly multiplexed assay to monitor pathogenicity, fungicide resistance and gene flow in the fungal wheat pathogen *Zymoseptoria tritici*"

### Supplementary Figure 1

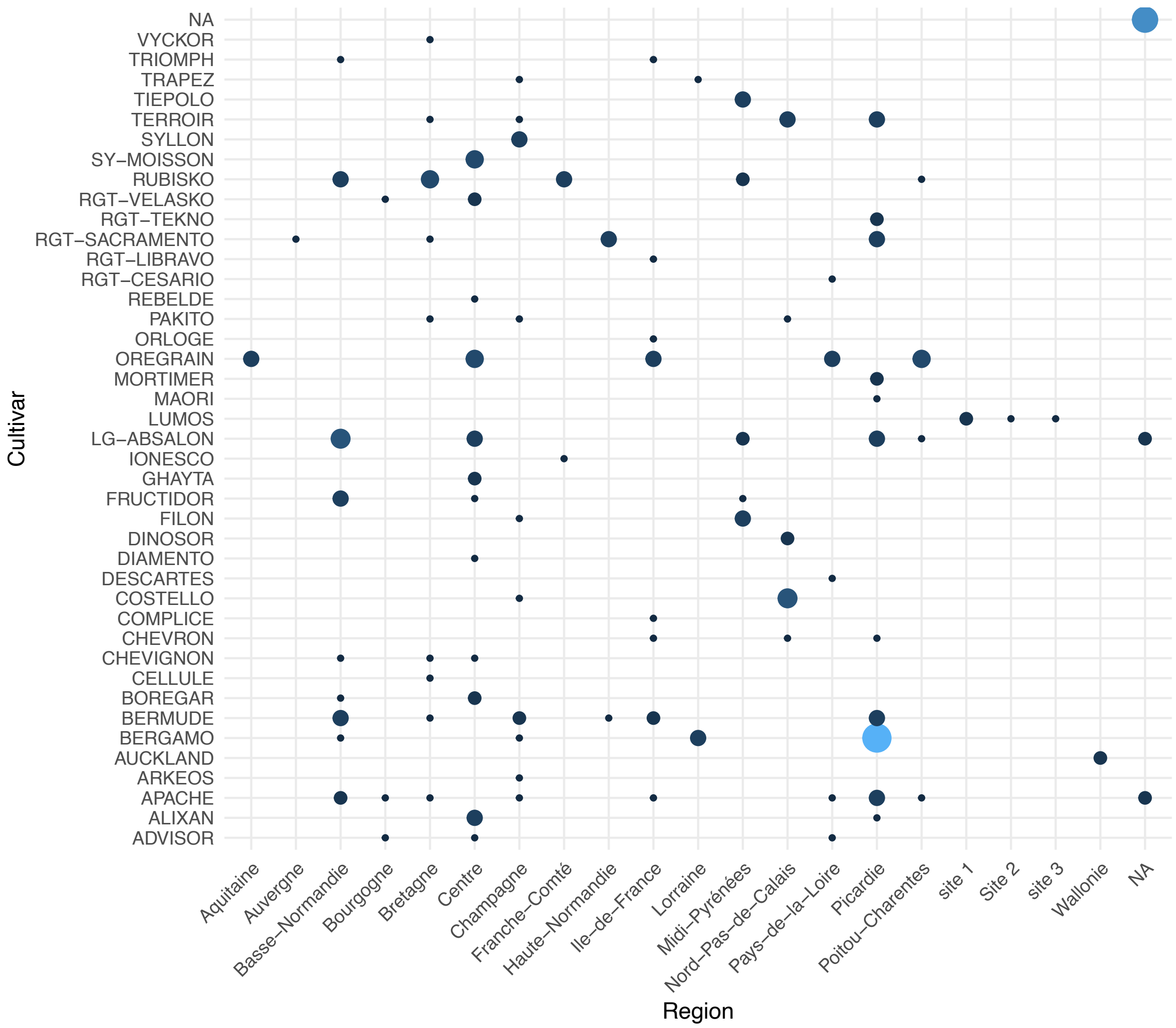


**Figure 1:** Wheat leaf samples collected in France, Belgium, Ireland and the United Kingdom separated by the cultivar of origin or unknown cultivar ("NA"). See File S1 (Table A) for details on the sample origins.
